# *In vitro* bio-activity of three potential antagonistic fungi against selected plant pathogens in Ghana

**DOI:** 10.1101/2024.08.07.606921

**Authors:** Elvis Dwamena, Richard Tuyee Awuah, Daniel Agbetiameh, Atta Kwesi Aidoo

## Abstract

In the quest of developing sustainable strategies for the management of diseases affecting major crop species, this study screened three antagonists, *viz. Trichoderma asperellum*., *Penicillium citrinum*. and *Myxotrichum stipitatum*., for inhibitory activity against seven plant pathogens, *viz. Pestalotia heterocornis, Curvularia eragrostidis, Lasiodiplodia theobromae, Sclerotium rolfsii, Colletotrichum gloeosporioides, Aspergillus flavus,* and *A. parasiticus* on Potato Dextrose Agar (PDA) plates using the dual culture method. For each potential antagonistic fungus-pathogen combination, four replicated dual culture plates were prepared and laid out in a completely randomized design (CRD). Control plates with only the pathogens and no antagonistic fungi were also established. All assay plates were incubated at 28 ℃ (approx.) and the diameters of the pathogen in both the dual culture and control plates were measured at 6, 10 and 14 days. Percentage inhibition of fungal pathogens were calculated and used to draw a colony reduction curve for each antagonist-pathogen interaction. Areas Under the Colony Reduction Curves (AUCRC’s) were calculated. Of the three antagonistic fungi tested, *Trichoderma asperellum*. was the most effective in inhibiting colony diameters of all seven pathogens with an average AUCRC of 813.19 % days, followed by the *Penicillium citrinum*. (519.25 % days). The weakest antagonist was *Myxotrichum stipitatum* sp. with an AUCRC of 440.6. % days. The results of the study suggest that *T*. *asperellum* and *P*. *citrinum* could be exploited for plant disease biocontrol.

## 1. Introduction

Biological control (bio control) agents (BCAs) are increasingly being utilized as alternative or complementary approach to synthetic chemicals in plant disease management because of the negative impacts of synthetic chemicals on the environment and human health[1]. Typically, biocontrol agents (BCAs) for plant diseases are fungi as well as bacteria and these organisms originate from the phyllosphere or rhizosphere and perform an essential role in the management of plant-pathogens[2].

The BCAs can demonstrate a variety of direct and indirect modes of action that achieve disease prevention. Examples of such mechanisms include antibiosis (where the antagonist produces an inhibitory metabolite or antibiotic against the pathogen); mycoparasitism (in which the antagonist obtains part or all of its nutrients from the pathogen); induced resistance (whereby a host plant defensive response against a plant pathogen is induced in the host plant) and growth enhancement (promotion plant growth while the effects of the disease is being reduced [3]. Many BCAs are now commercially available for use alone or in combination with chemicals against plant infections, thereby reducing the quantity of pesticides required. *Trichoderma* species are the most widely utilized fungal species in biological management of plant diseases [4]. *Trichoderma* species are non-sexual organisms that co-exist with other soil-dwelling fungi with teleomorphs of the Hypocrea genus, Ascomycota [5]. A variety of *Penicillium* species have been identified as antagonists to causative agents of plant diseases with methods of action such as induced resistance [6], the synthesis of antimicrobial compounds, as well as the development of mycoparasitic contacts [7],[8]. Studies have shown that the culture filtrate of *P. janczewskii* showed some inhibitory activity towards *Rhizoctonia solani* [7].

In Ghana, little progress has been made in the discipline of biocontrol technology against major plant pathogens, with a few studies reported by [9], [10], [11], [12], and [13]. Fungal and bacterial BCAs are promising for use because they are target-specific, have a rapid rate of reproduction, and short generation times [14]. Pathogens are exposed to pesticide resistance early due to increased pesticide usage. This makes disease control with chemicals less effective in the long run [15].

*Myxotrichum stipitatum* is a type of fungus that belongs to the *Myxotrichaceae* family.

*M*. *stipitatum* is associated with mycorrhizal interactions and an extensive search of the literature on the antifungal properties indicates that the fungus *M. stipitatum* has not been extensively studied.

It is imperative to periodically screen microorganisms for activity towards plant pathogens with a view to developing promising microorganisms into biocontrol agents. The research therefore, sought to periodically screen microorganisms for activity towards plant pathogens with a view to developing promising microorganisms into biocontrol agents, thus search for fungi with biocontrol potentials against seven plant pathogenic fungi for eco-friendly management option.

## 2. Materials and Methods

### 2.1 Sources of plant pathogens used in the study

The sources of the fungi used in the study are indicted in Table 1.

**TABLE 1:**
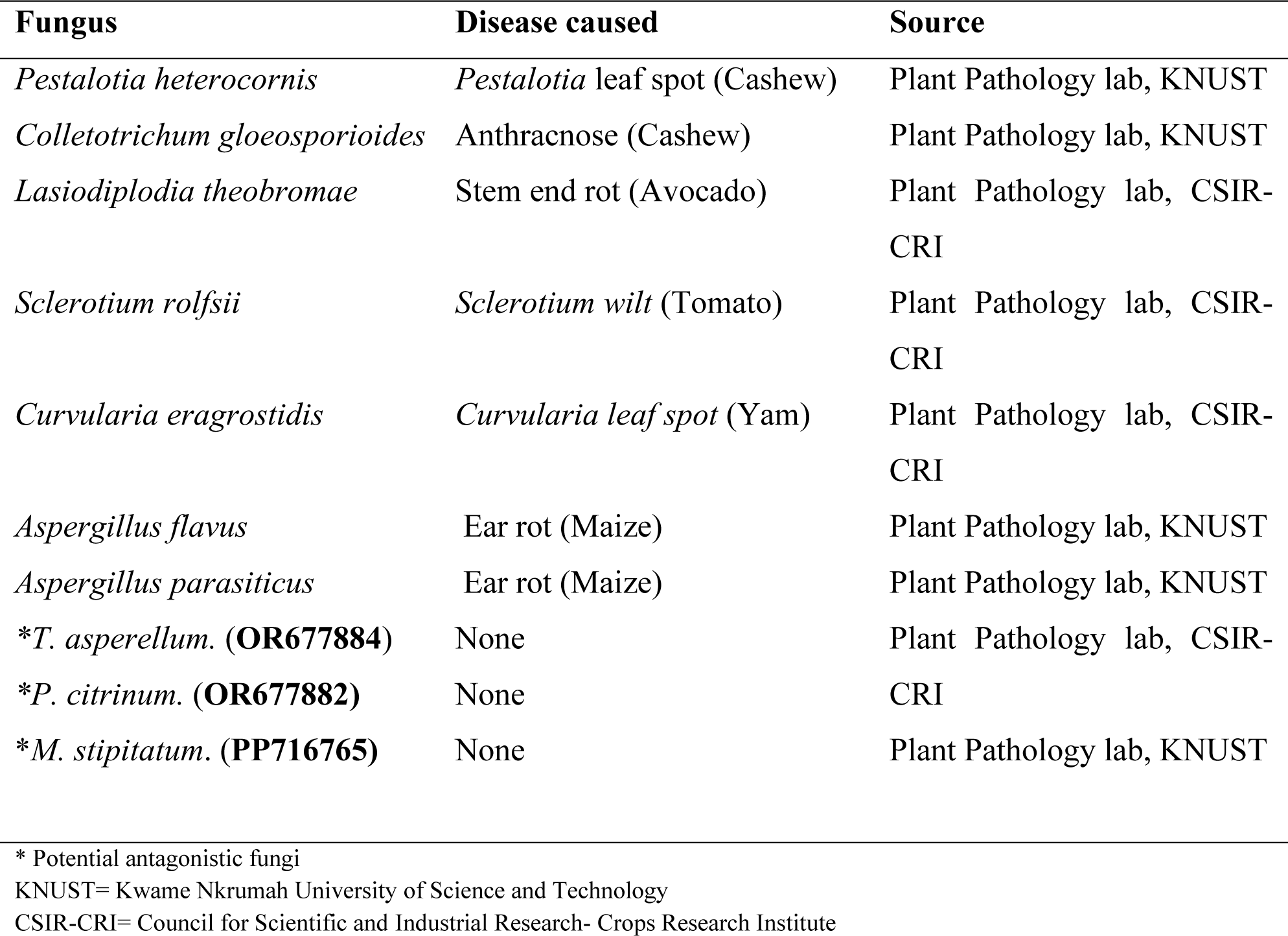
Test plant pathogens, fungal antagonist and their sources.

### 2.2 Test for fungal antagonism

The assay for antagonism was performed on artificial PDA using the dual culture technique described by [16]. A PDA-containing plate was seeded on one side (2 cm away from the margin of the plate) with a 5-mm-diameter mycelial plug obtained from the margin of a seven-day-old culture of the *Trichoderma* isolate. On the opposite side, a distance of 5 cm away from the potential antagonistic *Trichoderma* sp., a 5 mm mycelial plug of the pathogen was placed. Mycelial plugs of the pathogenic fungi were placed at one side at a distance of 2 cm away from the margin to establish a control plate. For *Penicillium* and the *Myxotrichum* isolates, mycelial bits were placed in 10 ml of sterile distilled water and manually shaken to dislodge the conidia into the water. For each of them, one end of a sterilized glass rod (0.5-cm-diameter) was vertically dipped into the spore suspension to pick the spores and tapped 2 cm from the margin of the test plate as done for the *Trichoderma* isolate. For each potential antagonistic fungus - pathogen combination, four replicate dual culture plates were prepared.

All assay plates were incubated at 28 ℃ (approx.) with approximately 8 h photoperiod of fluorescent white light. Diameters of the pathogen in both the dual culture and control plates were measured exactly on the 6^th^, 10^th^, and 14^th^ day after seeding. Percent inhibition of mycelial growth of the test fungi by the antagonistic fungi was calculated with the formula:

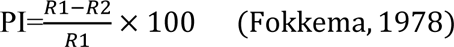

where,

PI = Percentage of inhibition of the pathogen;

R1 = Diameter of the pathogen in the control plate

R2 = Diameter of the pathogen in the dual culture plate

The percent inhibition data were then used to draw graphs of growth inhibition with time. Additionally, interaction between the three potential antagonistic fungi was also scored using the rating scale of [17] as shown in Table 2. This work was repeated once as a validation of the obtained results, three months after the completion of the first experiment.

**TABLE 2:**
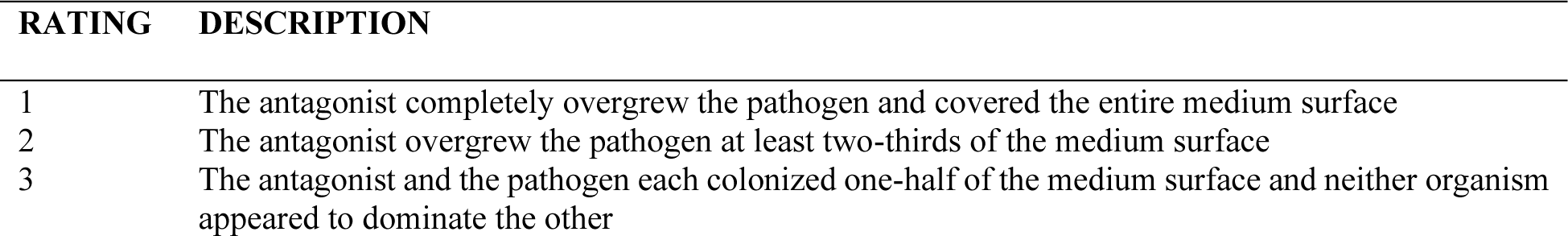
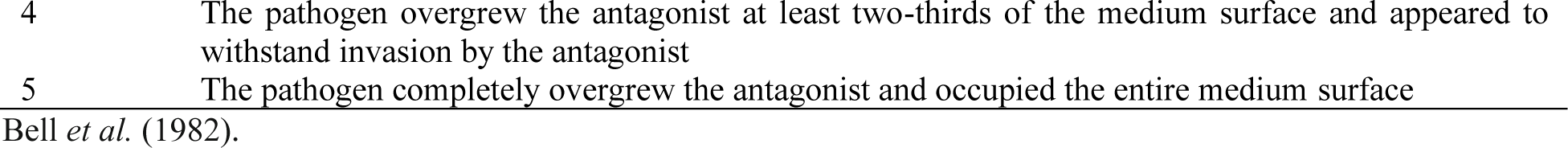
Rating scale of interaction between antagonistic fungi and pathogens.

### 2.3 Experimental Design and Data Analysis

Experimental design was Complete Randomised Design (CRD) factorial with two factors (Test plant pathogen and antagonistic fungus). Data sets for both the main and the repeated experiments were pooled together using the average pooling method before the analysis. Normality testing was conducted using Shapiro-Wilk test to evaluate whether our sample data follows the normal distribution. It revealed that the data did not significantly deviate from normality (P>0.05), thus, it supports the normality assumption for subsequent parametric tests. Data for AUCRCs were arcsine transformed and subjected to Analysis of variance (ANOVA) with Statistix software 9.0 [18]. The means of AUCRCs were separated with Tukey’s HSD at a significance level of 1%.

## 3. RESULTS

### 3.1 Test for fungal antagonism

Antagonism between *T. asperellum*, *P. citrinum* and *M. stipitatum* are shown in Figure 1.1, 1.2, and 1.3 respectively.

**FIGURE 1.1:**
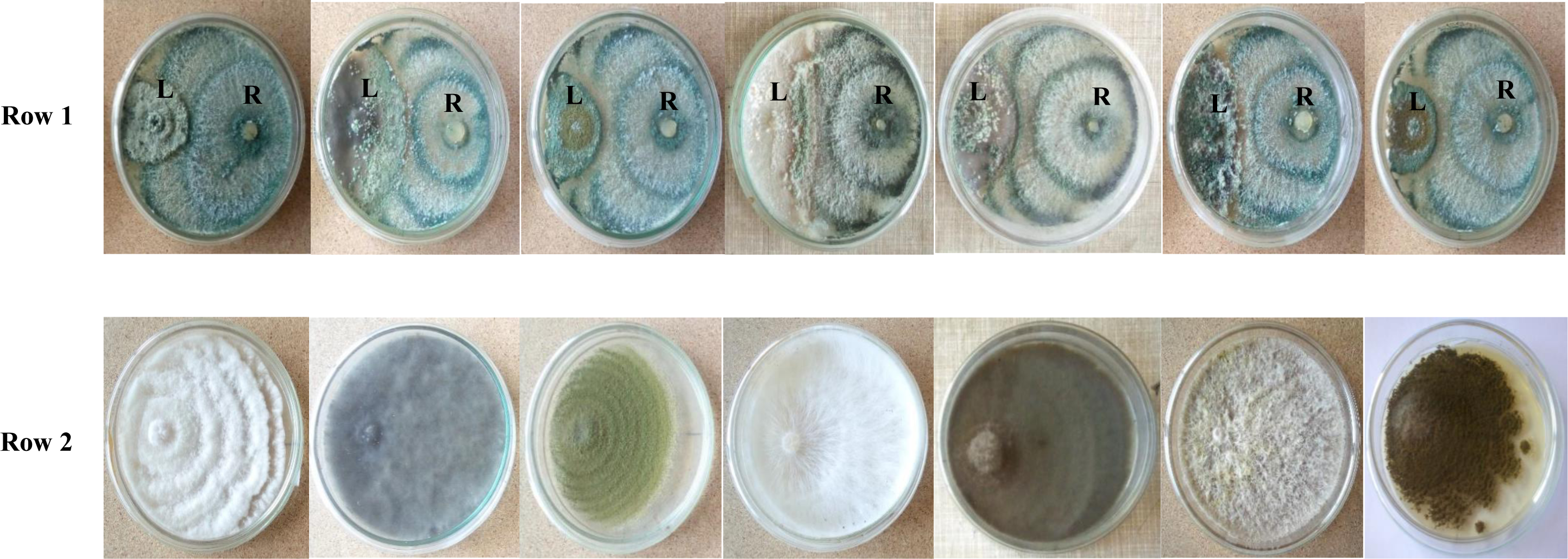
Pairings between *Trichoderma asperellum* and some plant pathogens. Row 1, obverse of cultures. On each plate, the antagonistic Trichoderma colony is on the right (marked R) and the plant pathogen colony is on the left (marked L). Row 2, obverse view of the coresponding control plates of the pathogens namely, (L-R) *Pestalotia heterocornis*, *Lasiodiplodia theobromae, Aspergillus flavus, Sclerotium rolfsii, Curvularia eragrostidis, Colletotrichum gloeosporioides,* and *Aspergillus parasiticus.* Cultures are 14 days old.

**FIGURE 1.2:**
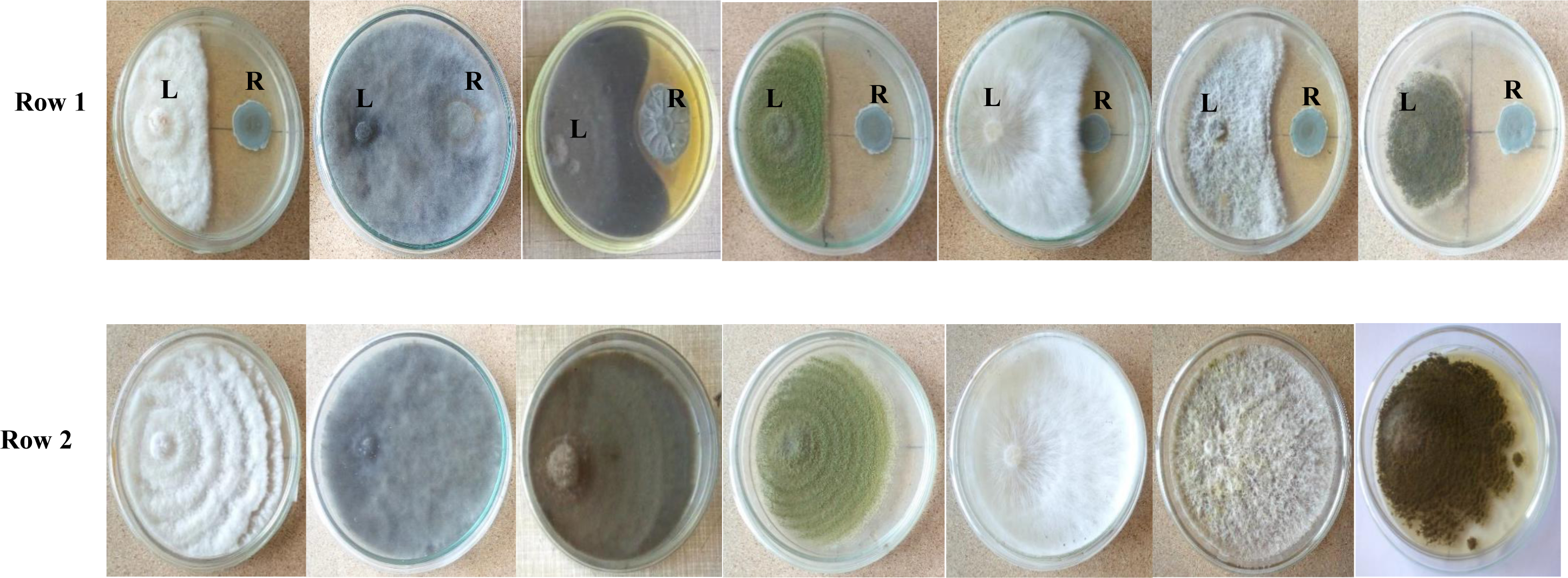
Pairings between *Penicillium citrinum* and some plant pathogens. Row 1, obverse of cultures. On each plate, the antagonistic *Penicillium* colony is on the right (marked R) and the plant pathogen colony is on the left (marked L). Row 2, obverse view of the corresponding control plates of the pathogens namely, (L-R) *Pestalotia heterocornis*, *Lasiodiplodia theobromae, Aspergillus flavus, Sclerotium rolfsii, Curvularia eragrostidis, Colletotrichum gloeosporioides* and *Aspergillus parasiticus.* Cultures are 14 days old.

**FIGURE 1.3:**
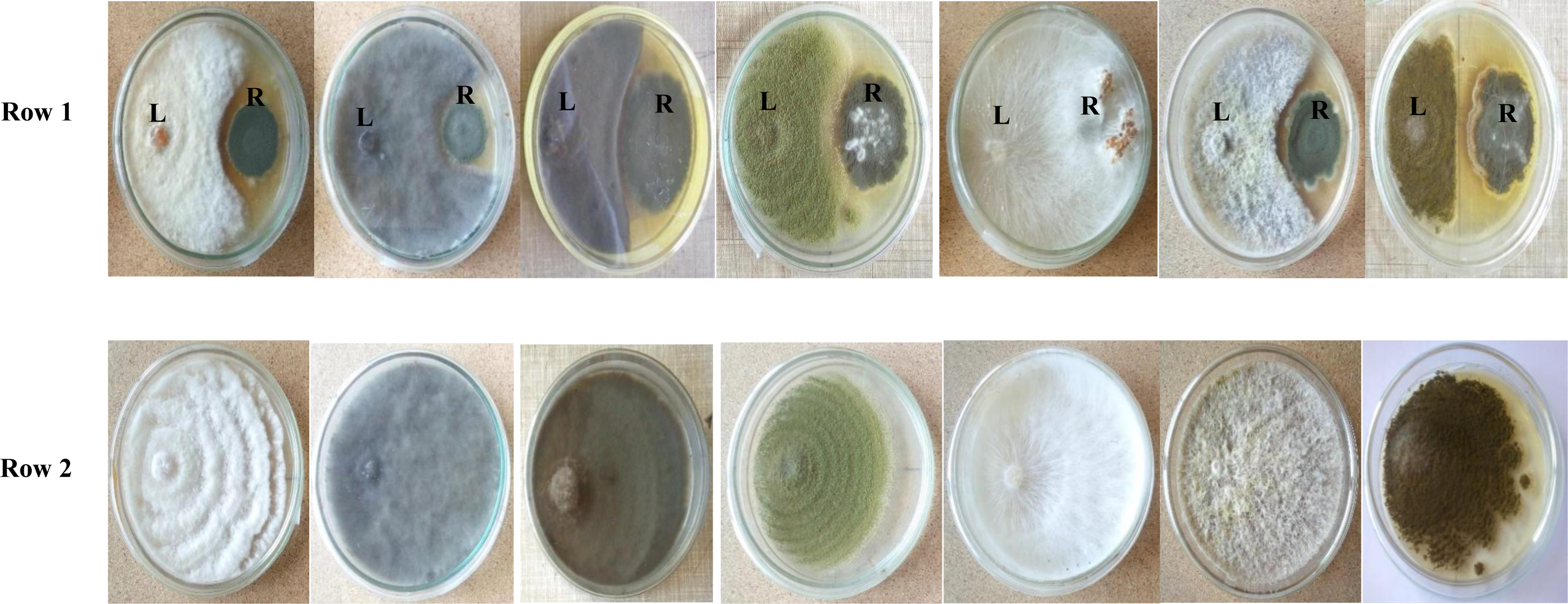
Pairings between *Myxotrichum stipitatum* and some plant pathogens. Row 1, obverse of cultures. On each plate, the antagonistic *Myxotrichum* colony is on the right (marked R) and the plant pathogen colony is on the left (marked L). Row 2, obverse view of the corresponding control plates of the pathogens namely, (L-R) *Pestalotia heterocornis, Lasiodiplodia theobromae, Aspergillus flavus, Sclerotium rolfsii, Curvularia eragrostidis, Colletotrichum gloeosporioides and Aspergillus parasiticus.* Cultures are 14 days old.

The antagonistic activity between *T. asperellum* and the seven plant pathogens shows the inhibitory potential of *T*. *asperellum* completely overgrowing the plant pathogens and covering the entire medium surface as well as overgrowing the pathogen at least two-thirds of the medium surface. Also, the inhibitory potential of *P. citrinum* against the seven plant pathogens clearly shows the formation of inhibition zones on almost all dual culture plates signifying the mechanisms of antibiosis. The inhibitory potential of *M. stipitatum* also shows some form of inhibition zones signifying the production of some metabolites by the antagonistic fungus.

*Trichoderma asperellum* had the highest AUCRC with an average of 813.19 % days against all the seven pathogens and inhibiting more than 50% of all the seven pathogenic fungi. The least effective of the three antagonist was *M. stipitatum* which had an average AUCRC of 440.60 % days (Table 3).

**TABLE 3:**
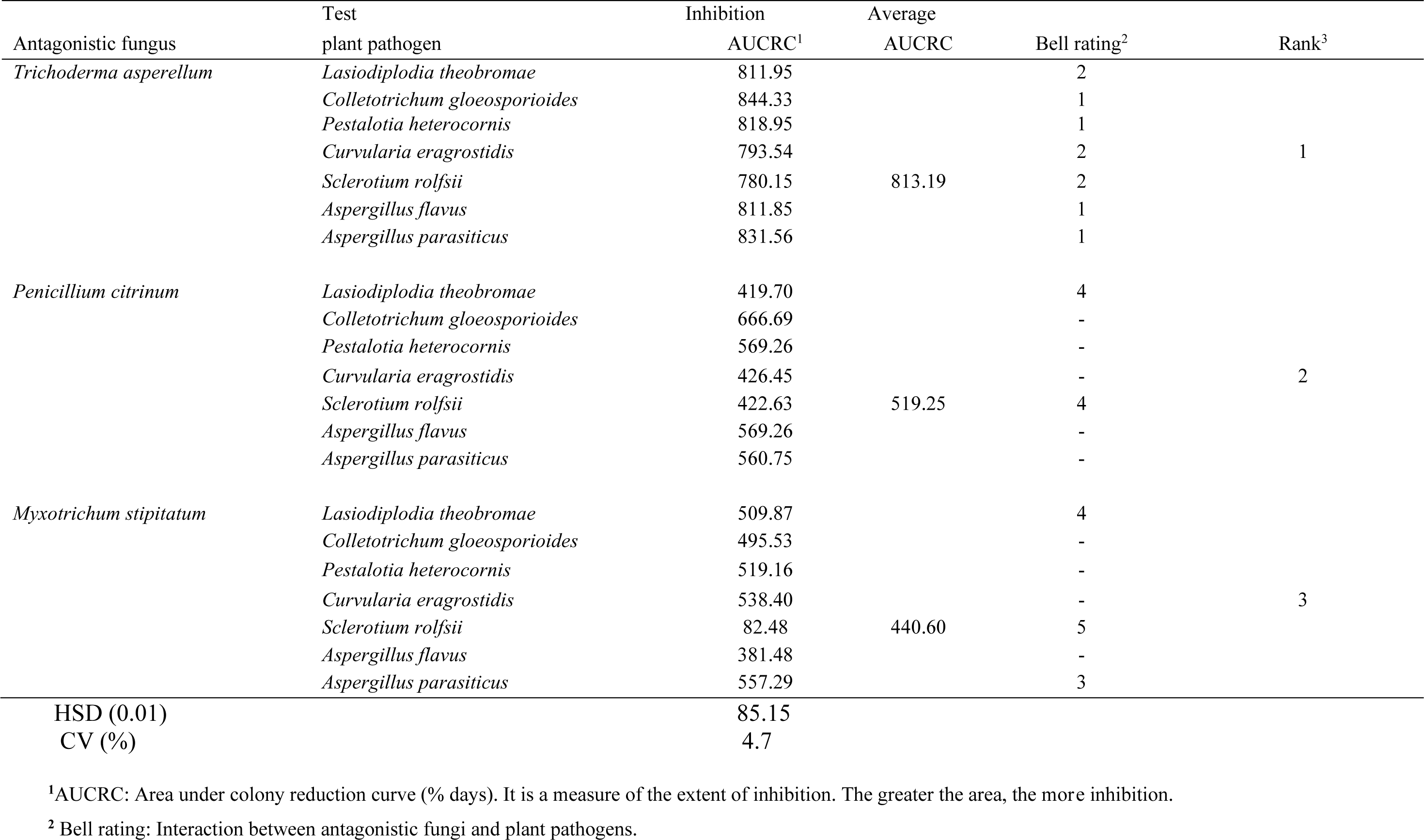
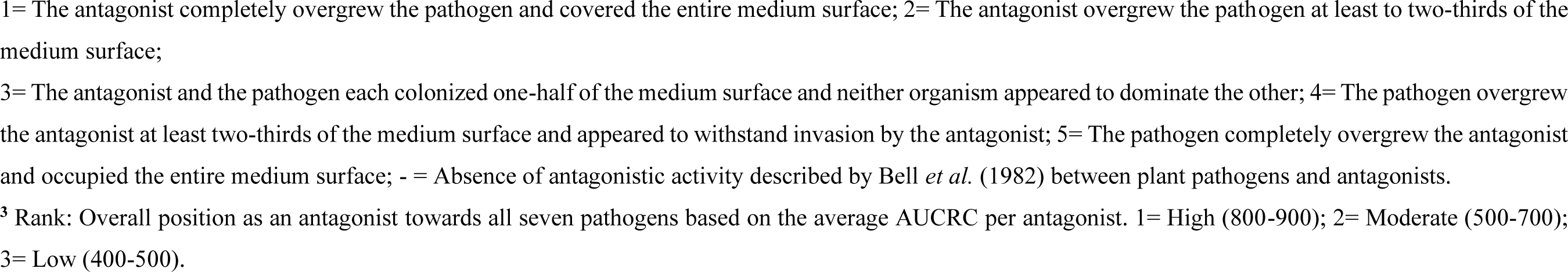
*In vitro* inhibition of the plant pathogenic fungi by three antagonistic fungi in the study.

| Antagonistic fungus | Test plant pathogen | Inhibition AUCRC<sup>1</sup> | Average AUCRC | Bell rating<sup>2</sup> | Rank<sup>3</sup> |
| --- | --- | --- | --- | --- | --- |
| *Trichoderma asperellum* | *Lasiodiplodia theobromae* | 811.95 | 813.19 | 2 | 1 |
|  | *Colletotrichum gloeosporioides* | 844.33 |  | 1 |  |
|  | *Pestalotia heterocornis* | 818.95 |  | 1 |  |
|  | *Curvularia eragrostidis* | 793.54 |  | 2 |  |
|  | *Sclerotium rolfsii* | 780.15 |  | 2 |  |
|  | *Aspergillus flavus* | 811.85 |  | 1 |  |
|  | *Aspergillus parasiticus* | 831.56 |  | 1 |  |
| *Penicillium citrinum* | *Lasiodiplodia theobromae* | 419.70 | 519.25 | 4 | 2 |
|  | *Colletotrichum gloeosporioides* | 666.69 |  | - |  |
|  | *Pestalotia heterocornis* | 569.26 |  | - |  |
|  | *Curvularia eragrostidis* | 426.45 |  | - |  |
|  | *Sclerotium rolfsii* | 422.63 |  | 4 |  |
|  | *Aspergillus flavus* | 569.26 |  | - |  |
|  | *Aspergillus parasiticus* | 560.75 |  | - |  |
| *Myxotrichum stipitatum* | *Lasiodiplodia theobromae* | 509.87 | 440.60 | 4 | 3 |
|  | *Colletotrichum gloeosporioides* | 495.53 |  | - |  |
|  | *Pestalotia heterocornis* | 519.16 |  | - |  |
|  | *Curvularia eragrostidis* | 538.40 |  | - |  |
|  | *Sclerotium rolfsii* | 82.48 |  | 5 |  |
|  | *Aspergillus flavus* | 381.48 |  | - |  |
|  | *Aspergillus parasiticus* | 557.29 |  | 3 |  |

| Antagonistic fungus | Test plant pathogen | Inhibition AUCRC<sup>1</sup> | Average AUCRC | Bell rating<sup>2</sup> | Rank<sup>3</sup> |
| --- | --- | --- | --- | --- | --- |
| *Trichoderma asperellum* | *Lasiodiplodia theobromae* | 811.95 | 813.19 | 2 | 1 |
|  | *Colletotrichum gloeosporioides* | 844.33 |  | 1 |  |
|  | *Pestalotia heterocornis* | 818.95 |  | 1 |  |
|  | *Curvularia eragrostidis* | 793.54 |  | 2 |  |
|  | *Sclerotium rolfsii* | 780.15 |  | 2 |  |
|  | *Aspergillus flavus* | 811.85 |  | 1 |  |
|  | *Aspergillus parasiticus* | 831.56 |  | 1 |  |
| *Penicillium citrinum* | *Lasiodiplodia theobromae* | 419.70 | 519.25 | 4 | 2 |
|  | *Colletotrichum gloeosporioides* | 666.69 |  | - |  |
|  | *Pestalotia heterocornis* | 569.26 |  | - |  |
|  | *Curvularia eragrostidis* | 426.45 |  | - |  |
|  | *Sclerotium rolfsii* | 422.63 |  | 4 |  |
|  | *Aspergillus flavus* | 569.26 |  | - |  |
|  | *Aspergillus parasiticus* | 560.75 |  | - |  |
| *Myxotrichum stipitatum* | *Lasiodiplodia theobromae* | 509.87 | 440.60 | 4 | 3 |
|  | *Colletotrichum gloeosporioides* | 495.53 |  | - |  |
|  | *Pestalotia heterocornis* | 519.16 |  | - |  |
|  | *Curvularia eragrostidis* | 538.40 |  | - |  |
|  | *Sclerotium rolfsii* | 82.48 |  | 5 |  |
|  | *Aspergillus flavus* | 381.48 |  | - |  |
|  | *Aspergillus parasiticus* | 557.29 |  | 3 |  |
| HSD (0.01) |  | 85.15 |  |  |  |
| CV (%) |  | 4.7 |  |  |  |

<sup>1</sup>AUCRC: Area under colony reduction curve (% days). It is a measure of the extent of inhibition. The greater the area, the more inhibition.
<sup>2</sup> Bell rating: Interaction between antagonistic fungi and plant pathogens.
1= The antagonist completely overgrew the pathogen and covered the entire medium surface; 2= The antagonist overgrew the pathogen at least to two-thirds of the medium surface;
3= The antagonist and the pathogen each colonized one-half of the medium surface and neither organism appeared to dominate the other; 4= The pathogen overgrew the antagonist at least two-thirds of the medium surface and appeared to withstand invasion by the antagonist; 5= The pathogen completely overgrew the antagonist and occupied the entire medium surface; - = Absence of antagonistic activity described by Bell *et al.* (1982) between plant pathogens and antagonists.
<sup>3</sup> Rank: Overall position as an antagonist towards all seven pathogens based on the average AUCRC per antagonist. 1= High (800-900); 2= Moderate (500-700); 3= Low (400-500).

## 4. Discussion

The inhibitory potential of the three antagonistic fungi against the seven plant pathogenic fungi varied in culture. Comparatively, the inhibitory potential of *T. asperellum* against the mycelial growth of all seven pathogenic fungi was the highest indicated by the Area Under the Colony Reduction Curves (AUCRCs). The concept of Area Under the Disease Progress Curve (AUDPC) which may be used in assessing plant disease development over time is applied to the Area Under the Colony Reduction Curve (AUCRC) in the current study. The lower the value of the area, the lower the disease development. This would be analogous to the AUCRC over time meaning that the lower the area of the colony reduction curve the lower the inhibition by the antagonistic fungus. This approach may be used in assessing the effects of some factors over time. The antagonistic activity of *T. asperellum* against economically important plant pathogenic fungi is well documented. *T. asperellum* has been shown to exhibit strong antagonistic potential against *Fusarium verticillioides* and *Ustilago maydis* in maize [19], *Rhizoctonia solani* causing sheath blight disease of rice [20]; *Fusarium oxysporium* causing Tomato wilt [21] and *Pythium myriotylum*, the causative agent of cocoyam root rot disease [22]. Generally, *Trichoderma* species exhibit unique biocontrol activity and produce many volatile secondary metabolites including Aldehydes, Hydrogen cyanide, Ketones and Ethylene. These chemical compounds are important in controlling plant pathogens [23], [24]. A study cited as [25] showed the inhibitory effect of *T. asperellum* against *P. megakarya* was attributed to be a direct penetration in the sporocystes (sporangium), the coil around the *P. megakarya* hyphae and the formation of appressoria on the hyphae surface causing their destruction, and the substantial activity of hydrolytic enzymes such as laminarinase.

The degree of inhibition as expressed in the AUCRC for *P. citrinum* was relatively lower for all the seven plant pathogens. In a similar study, [26] screened five *penicillium* species including *P*. *citrinum* against *Macrophomina phaseolina* and reported that *P*. *citrinum* was the second most effective in suppressing the mycelia growth of the pathogen. Similarly, [27] reported that, *P*. *citrinum* culture filtrate at different concentrations, inhibited the germination and growth of Sclerotia of *Sclerotinia minor* [27]. Inhibition zones created by *P. citrinum* could have been due to the production of antifungal compounds such as citrinin and emodin [28]. These antifungal compounds have been demonstrated to be inhibitory towards some plant pathogens [29]. In the current study, antifungal compounds produced by *P*. *citrinum* could possibly be responsible for inhibition of mycelial growth of the seven plant pathogens evaluated in this study.

*Myxotrichum stipitatum* also possesses some antifungal properties, although it had the lowest AUCRC amongst the three antagonistic fungi. It formed inhibition zones against all plant pathogens tested with the exception of *S. rolfsii*, and this shows that it could produce some antifungal compounds into the medium resulting in the formation of inhibition zones against the pathogenic fungi. This biocontrol agent has not been extensively studied. The current study is the first report in Ghana especially on its antifungal potency. The only reason *M. stipitatum* was tested in the current study was because of its resemblance with *Metarhizium anisopliae* which also has antifungal properties [30].

## 5. Conclusion

When the three antagonistic fungi were tested against all the seven plant pathogens for biocontrol potential using the dual culture technique, *T. asperellum* exhibited the strongest antagonistic activity against all seven pathogens followed by *P. citrinum.* The least inhibitory antagonist was *M. stipitatum.* Observations from the dual culture set-ups of antagonists paired against plant pathogens suggest that *T. asperellum* and *P*. *citrinum* may be developed into biocontrol agents.

## Abbreviations

ANOVA: Analysis of Variance
BCAs: Biocontrol agents
CRD: Completely randomized design
CRI: Crops Research Institute
CSIR: Council for Scientific and Industrial Research
CV: Coefficient of Variation
HSD: Honesty Significant Difference
KNUST: Kwame Nkrumah University of Science and Technology
PDA: Potato Dextrose Agar

## Data Availability

The data are also available from the corresponding author upon request. Please contact Elvis Dwamena at for more information.

## Authors’ Contribution

E.D. played a role in conducting the experiment, collecting the data, and helped draft the manuscript. R.T.A. critically revised the thesis before writing of manuscript. D.A. critically revised the manuscript and A.K.A. analyzed the data and critically revised the manuscript. All authors gave their final approval and agreed to be accountable for all aspects of this work.

## Acknowledgment

Special appreciation and recognition to staff of Department of Crop and Soil Sciences, (Plant pathology laboratory) KNUST, CSIR-Crops Research Institute (Plant pathology laboratory), and Cocoa Research Institute of Ghana (Biotechnology Laboratory), Akim-Tafo.

